# Decoupling synaptic weight and connection sparsity reveals asymmetric control of network dynamics

**DOI:** 10.1101/2025.04.02.646896

**Authors:** Tea Tompos, Fleur Zeldenrust, Tansu Celikel

**Affiliations:** Donders Institute for Brain, Cognition and Behaviour, Radboud University, Nijmegen, The Netherlands; School of Psychology, Georgia Institute of Technology, Atlanta - GA, United States of America; Georgia Tech-CNRS IRL 2958, Georgia Institute of Technology - Europe, Metz, France

**Keywords:** connectivity probability, functional heterogeneity, network dynamics, spiking neural networks, synaptic weight

## Abstract

Circuit dynamics arise from the interaction between the network’s connectivity structure and intrinsic neuronal nonlinearities, yet the roles of key structural parameters — synaptic weight (W) and connection probability (P) — are usually examined in simplified network models. Using a biologically grounded, multilayer spiking model of thalamocortical microcircuitry, incorporating conductance-based neurons and rodent somatosensory cortex connectivity, we systematically scaled W and P and identified four organising principles of population dynamics. First, stronger synapses monotonically amplified spiking across all populations. Second, increasing connection density produced a weight-dependent bidirectional outcome: adding weak synapses preserved baseline activity, whereas adding strong ones suppressed firing. Third, concurrent increases in W and P yielded sublinear effects, where population activity increased less than expected from the sum of their individual impacts. Fourth, two functional neuronal classes emerged — scaling-invariant neurons that reliably transmitted thalamic input across connectivity regimes, and variant neurons that spiked selectively under specific connectivity scales. These classes differed in their excitation–inhibition balance, shaped by the strength of recurrent inhibition. Together, our findings show that synaptic weight and connection probability work in concert to define cortical operating regimes and generate the functional diversity in neuronal responses that supports flexible computation.

## 2. Introduction

The computational activity of neural circuits arises from both the intrinsic properties of individual neurons and the structure of their connections. Two fundamental synaptic parameters underpin this structure: synaptic weight (W), reflecting the strength of individual connections, and connection probability (P), determining the sparsity or density of network connectivity. Although their roles have been widely studied (Harris and Mrsic-Flogel 2013; Gao and Ganguli 2015; Shimoura et al. 2021), a deeper understanding of their influence on circuit-level dynamics — especially in realistic, nonlinear systems — remains limited. Most prior models rely on analytical tractability of mathematically simplified systems, often invoking scaling laws that conflate the two structural parameters (van Albada et al. 2015; Barral and D Reyes 2016; Brunel 2000; van Vreeswijk and Sompolinsky 1996; Golomb and Hansel 2000). When W and P were considered independently (Harris et al. 2023; Vegué et al. 2025; Mastrovito et al. 2024), analyses defining the roles remain limited to simplified systems, omitting the nonlinear dynamics characteristic of biological neurons.

Here, we investigate how W and P shape activity states in a multi-layer spiking neural network composed of Hodgkin–Huxley neurons. This framework enables us to ask what computational properties or dynamical activity states emerge when synaptic weights and connection sparsity are systematically dissociated in a more realistic setting.

Both synaptic weight and sparsity are central to theories of brain computation and learning. Even in simplified models, changing the mean or variance of synaptic weights can profoundly alter overall firing rates, generate chaotic dynamics, or drive transitions between synchronous and asynchronous firing regimes (Vegué et al. 2025; Mastrovito et al. 2024; Flesch et al. 2022; Brunel 2000). Changing the distribution shape — for instance, rendering synapses sparse but strong — can increase network responsiveness to weak inputs (Iyer et al. 2013; Kriener et al. 2014). Moreover, plastic changes in synaptic weights promote learning and memory formation in spiking networks (Tetzlaff et al. 2013; Auth et al. 2020; Bicknell and Latham 2025). Independent of weight distributions, sparse connectivity itself benefits computational efficiency, memory storage, and temporal processing compared to fully connected networks (Knoblauch et al. 2014; Knoblauch and Sommer 2016; Feng and Brunel 2022; Fruengel and Oberlaender 2025).

Despite these advances, the precise influence of W and P on neuron and network activity within realistic regimes remains unclear. The nature of neuronal nonlinearity is crucial: distinct coding schemes can arise from the same network structure under different neuron models. For example, van Meegen and Sompolinsky (2025) demonstrate that linear networks rely on analogue codes, whereas nonlinear networks with rectified or sigmoidal neurons adopt sparse or redundant coding schemes, respectively, even when achieving identical task accuracy. These findings motivate our focus on biologically grounded spiking models to reveal how connectivity governs emergent dynamics.

Using empirically constrained connectivity and realistic neuronal responses derived from the rodent somatosensory system, we established a biologically plausible activity baseline in a thalamocortical network model, then varied W and P independently to obtain a two-dimensional W-P map of network dynamics.

Our findings reveal an asymmetric influence: synaptic weight is a dominant driver of neuronal spiking, with global up- or down-scaling modulating firing rates more than changes in connection probability. In contrast, a shift from a sparsely to densely connected network via upscaling P exerts bidirectional effects that depend on synaptic weight. Adding weak connections leaves firing rate largely unchanged, whereas adding strong connections suppresses activity by recruiting widespread inhibition. When weight and probability were scaled together, sublinear interactions emerged across cortical layers, coinciding with increased inhibitory recruitment that potentially imposes an effective “ceiling” on activity. Additionally, we identify two functional subclasses of neurons: scaling-invariant neurons, which easily transition from balanced to excitation-dominated states as we scale connectivity parameters, maintaining stimulus responsiveness over a wide parameter range; and scaling-variant neurons, which exhibit selective recruitment depending on W-P scaling and maintain a tight E/I balance. The two classes are strongly separable based on the strength of recurrent inhibition that neurons receive.

These results indicate that exploring both synaptic structural parameters in realistic models uncovers dynamics far richer than predicted by simplified frameworks. Independent tuning of weight and sparsity can jointly regulate global activity, shape interlayer communication, and drive functional heterogeneity, illuminating how microscopic connectivity underpins macro-scale computation and adaptation.

## 3. Methods

### 3.1. Model Architecture

#### 3.1.1. Thalamocortical Network Design

The spiking neural network (SNN) model consisted of a thalamic input layer and two cortical layers (L), reflecting the laminar organisation in rodent primary somatosensory cortex (S1). The cortical layers L4 and L2/3 were downscaled compared to the *in vivo* microcircuit, with neuron counts scaled to 5% of those in a typical rat cortical column (221 neurons in L4, 287 in L2/3) (Meyer et al. 2010). Within each layer, neurons were assigned as excitatory (E, 85%) or inhibitory (I, 15%), yielding populations of L4E (188), L4I (33), L2/3E (244), and L2/3I (43). The full SNN comprised 13 distinct projection types, each defined by empirical synaptic weight (W) and connection probability (P) parameters (Markram et al. 2015), such that, for each population pair (e.g., L4E → L4I), P determined the proportion of realized synapses (e.g., 4.2% for L4E → L4I), each assigned an identical W. All baseline W and P values are listed in Table 3.

#### 3.1.2. Thalamic Layer as Input

The thalamic input layer comprised two subpopulations with arbitrary sizes: 45 pattern-generating and 24 background thalamocortical relay neurons. Pattern-generating population was designed to fire synchronously in brief windows, delivering event-based input to L4 (Supplementary Figure 1b-d). The background group spiked asynchronously to simulate thalamic noise. The two spiking regimes (i.e. synchronous and asynchronous) were controlled by the neuron-specific external current *I*_*ext*_ (Eq. 3), where random spike times were distributed based on a Poisson process. Pattern generators received a shared *I*_*ext*_ template, but with temporally jittered spike times for each neuron (jitter times drawn from a Gaussian distribution: mean 0 ms, standard deviation (SD) 5 ms); background neurons received unique *I*_*ext*_ templates with uncorrelated spike times. All *I*_*ext*_ templates were convolved with a double-exponential kernel to approximate naturalistic excitatory postsynaptic potential (EPSP) kinetics (Eq. 3.2.1; Supplementary Materials).

### 3.2. Single Neuron Models

#### 3.2.1. Cortical Neurons

All cortical neurons were modelled as single compartments governed by the Hodgkin-Huxley formalism:

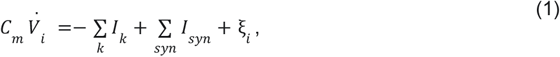

where *V*_*i*_ is membrane potential, *C*_*m*_ is membrane capacitance, *I*_*k*_ are voltage-gated channel currents, *I*_*syn*_ are receptor-mediated synaptic currents, and ξ_*i*_ is additive Gaussian noise (mean 0 pA, SD 100 pA, updated at 0.01 ms). Four channels were included, where *k* in *I*_*k*_ stands for: sodium (Na), delayed-rectifier potassium (KDR), slow non-inactivating potassium (M), and leak (L). Channel kinetics followed minimal Hodgkin-Huxley models for rodent S1 E and I neurons (Pospischil et al. 2008):

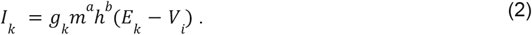

where *g*_*k*_ is maximal conductance, *E*_*k*_ is reversal potential, *m*^*a*^ defines activation and *h*^*b*^ inactivation dynamics with channel-specific exponents (see Supplementary Materials, Eqs. 2.1–2.4; parameters in Supplementary Table 1).

##### Model fitting

Template model parameters (Pospischil et al. 2008) were adjusted to fit empirical frequency–current (I–f) curves for S1 E and I neurons (Zeldenrust et al. 2024) to achieve a closer match between empirical and modelled neurons (Supplementary Figure 1a and Supplementary Table 1).

#### 3.2.2. Thalamic Neurons

Excitatory thalamocortical relay (TCR) neurons were also single-compartment Hodgkin-Huxley models, governed by:

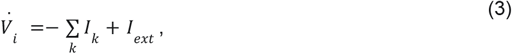

where *I*_*ext*_ is the external input current for triggering neuron-specific activity (Eq. 3.2, Supplementary Materials). Following the TCR model from (Ching et al. 2010), *k* in *I*_*k*_ stands for: sodium (Na), potassium (K), T-type calcium, hyperpolarisation-activated (h), potassium leak (KL), and general leak (L) current (Eqs. 3.1.1 — 3.1.6, Supplementary Materials).

##### Model Fitting

The TCR model from (Ching et al. 2010) was developed to simulate the rodent ventral posteromedial (VPM) thalamic nucleus during anaesthesia. To mimic VPM spiking in awake states (Urbain et al. 2015), resting membrane potentials were depolarised by increasing h-current conductance (*g*_*h*_), and the amplitude of external drive was increased in pattern-generating and background TCRs, with Poisson input rates reduced (parameter values in Supplementary Table 2).

### 3.3. Synaptic Transmission

#### 3.3.1. Synapse Model

Cortical neurons received AMPA (excitatory) and GABAa (inhibitory) synaptic currents:

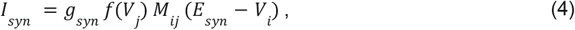

where *syn* stands for AMPA or GABAa, *g*_*syn*_ is maximal synaptic conductance, *E*_*syn*_ is reversal potential, *f*(*V*_*j*_) is a presynaptic voltage-dependent receptor activation (Eq. 4.1, Supplementary Materials), and *M*_*ij*_ represents pre (*i*) and post (*j*) population connectivity. Parameters are detailed in Supplementary Table 3.

#### 3.3.2. Synaptic Scaling

To normalise E and I currents before imposing synapse-specific weights, we scaled AMPA and GABAa conductances (*g*_*syn*_ in Eq. 4) to match the amplitudes of unitary excitatory and inhibitory postsynaptic potentials (uEPSP and uIPSP, respectively) evoked by one presynaptic action potential. The range of uEPSP and uIPSP amplitudes for synapses in cortical L4 and L2/3 is between 0.5 and 7 mV (Markram et al. 2015). We matched uEPSP and uIPSP amplitudes at 2.3 mV, which is within the expected range. For thalamocortical contacts, *g*_*AMPA*_ was doubled compared to corticocortical *g*_*AMPA*_ to compensate for the population size mismatch between VPM and L4, ensuring effective cortical recruitment by thalamic input (Supplementary Table 3).

#### 3.3.3. Connectivity Matrices

Synaptic connectivity between presynaptic (*j*) and postsynaptic (*i*) populations was implemented using projection-specific connectivity matrices. Each matrix *M*_*ij*_ (Eq. 4) was parameterised by *W*_*ij*_ and *P*_*ij*_ (Table 3). For each projection type listed in Table 3, the total number of synapses 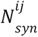 was calculated by multiplying the presynaptic population size (*N*_*j*_), the postsynaptic population size (*N*_*j*_), and the connection probability (*P*_*ij*_). We initialised each connectivity matrix *M*_*ij*_ as a sparse binary matrix with a randomly distributed 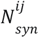 number of active synapses. Synaptic weights were then applied via element-wise multiplication: *M*_*ij*_ ⊙ *W*_*ij*_ (Figure 1c-d).

**Figure 1:**
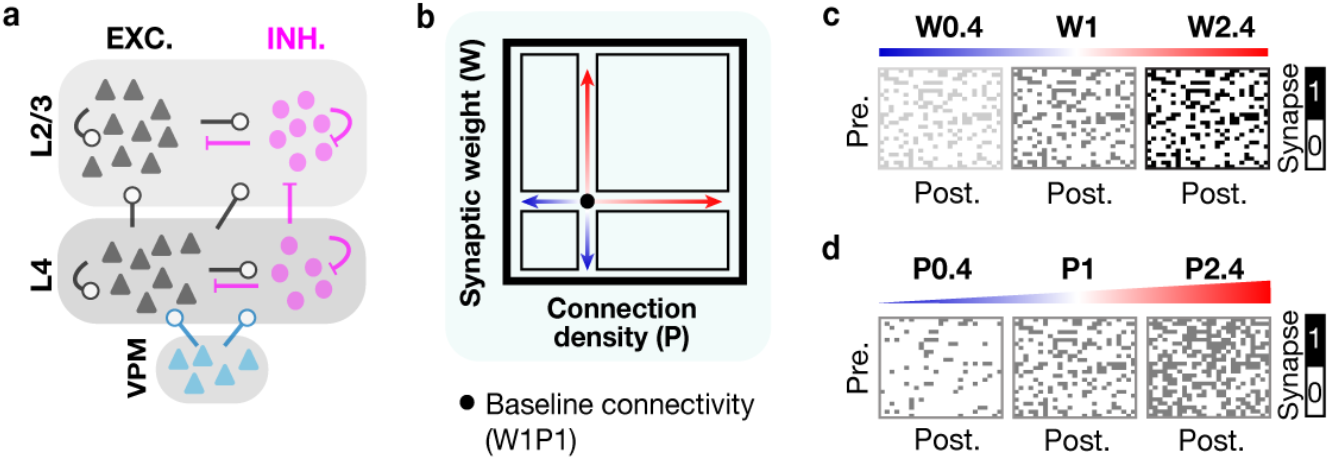
Studying the effects of synaptic weight (W) and density (P) scaling in a multi-layer spiking neural network. (a) The network architecture: Cortical layers (L) 4 and 2/3 contain excitatory (EXC., black triangles) and inhibitory (INH., pink circles) neurons. Layers have recurrent (within layers) and feedforward (between layers) connections (edges with circle tips, excitatory connections; pink edges with line tips, inhibitory connections). Both L4 populations receive spiking activity from excitatory neurons (blue triangles) in the thalamic ventral posteromedial (VPM) layer. Weights and neuron numbers are set according to Table 3 (Methods). (b) A cartoon of the experiment — a scaling of weight (W) and connection density via probability (P) in a 2D parameter space with respect to the reference values W1 and P1 (black dot). Red arrow part, increase in W or P; blue arrow part, decrease in W or P. (c) Example realisation of one connectivity matrix for a random connection edge from a. Upcaling W from the baseline (W1→W2.4) increases the strengths of all existing synapses, while downscaling (W1→W0.4) decreases their strength. Pre., presynaptic population; Post., postsynaptic population. Black/grey dots are existing synapses. White space means no synapse. (d) Same as c but for P instead of W. Upscaling P from baseline (P1→P2.4) randomly adds new synapses with identical strength, while downscaling P (P1→P0.4) randomly eliminates synapses.

### 3.4. Connectivity scaling simulations

#### 3.4.1. Parameter Space Design

To model connectivity changes in the network, we systematically scaled W and P across a symmetric 2D parameter space with a shared vector of scaling factors (Figure 1b). Scaling factors ranged from 0.4 to 2.4 in 0.2 increments. For each scaling condition (WxPy), x scaled W and y scaled P relative to their connection-specific baseline values (Table 3). Baseline values were unscaled in the W1P1 condition, which represented the initial network connectivity state. Note that scaling modified all cortico-cortical projections across the network (i.e., all connectivity matrices *M*_*ij*_), while thalamocortical projections were not scaled.

#### 3.4.2. Simulation Protocol

Simulations were executed in MATLAB using the DynaSim toolbox (Sherfey et al. 2018). Each network realisation ran for 800 ms (dt = 0.01 ms), integrating with a fourth-order Runge-Kutta algorithm. The first 100 ms of activity were excluded from analysis to avoid initial transients. For each unique W-P scaling condition, 10 network realisations were performed. All were executed using a fixed random seed for cortical network structure and noise, ensuring identical network architectures and noise traces within and across conditions for direct comparability. Thalamic spiking, however, was independently resampled for each network realisation using a seed-independent Poisson process. This resulted in naturalistic trial-to-trial variability in cortical responses for every W-P scaling condition, without confounding changes in internal circuit structure or noise.

## 4. Results

We built a reduced model of the canonical thalamocortical microcircuit, consisting of a ventral posteromedial (VPM) thalamic input layer (L) and two cortical layers connected in a feedforward fashion: VPM→L4→L2/3 (Figure 1a, Methods). Each projection used layer- and cell-type–specific synaptic weight (W) and connection probability (P) (Table 3). Transient thalamic synchrony elicited realistic spiking in L4 and L2/3, matching in vivo S1 population responses (Supplementary Figure 1b–e). From the baseline W1P1 configuration, we scaled W and P across a two-dimensional grid (Figure 1b), covering W > 1 (synaptic potentiation) or W < 1 (depression) and P > 1 (synaptic growth) or P < 1 (elimination), where 1 is “unscaled” (Figure 1c–d). Following connectivity scaling, we performed a comprehensive analysis of the resulting networks presented below.

### 4.1. Synaptic Weight Scaling Dominates Over Connection Density in Gating Population Activity

We first examined how L4 population activity – defined as the total evoked spikes (S) per E or I populations (Supplementary Materials) – varies with W and P scaling relative to baseline. Average spike outputs mapped across W-P space showed that high W/low P gave maximal activity, while low W/high P showed minimal output for both L4E and L4I (Figure 2a). These distributions were asymmetric relative to the WxPy antidiagonal, where *x* and *y* scaling factors are equal. Specifically, positive ΔS = S(WxPy) – S(WyPx) highlighted a dominant effect of W over P (Supplementary Figure 2a). Furthermore, increasing P did not increase spiking when W<1, and at high W, further increases in P contributed little to output. This identifies synaptic weight as the primary determinant of L4 activity.

**Figure 2:**
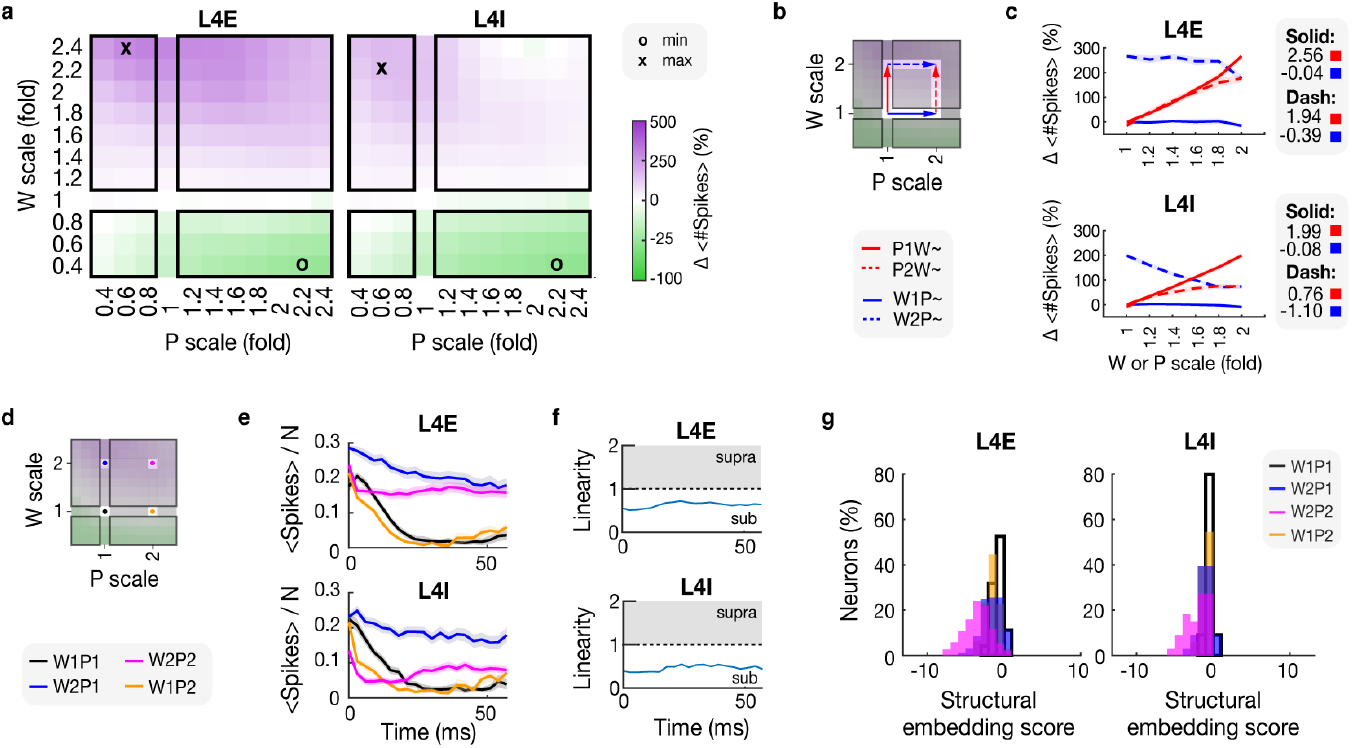
Modulation of spiking activity in the input layer 4 by changes in synaptic weights and density. (a) Heatmaps for L4 excitatory (E, left) and L4 inhibitory (I, right) show the relative change (%) in the mean spike number (#) with respect to W1P1 (colour scale on the right; shared for both maps). Synaptic weights (W, vertical axis) and density (P, horizontal axis) were scaled symmetrically (scaling factors: 0.4-2.4, increments of 0.2). Symbols on maps point to conditions with the maximal positive change (x) and the maximal negative change (o). (b) A map (top) illustrates the four analysed W-P transition pathways (see legend below): increasing W at baseline (P1W∼, solid red) or doubled P (P2W∼, dashed red), and increasing P at baseline (W1P∼, solid blue) or doubled W (W2P∼, dashed blue). (c) Line plots (top, L4E; bottom, L4I) show a relative change (%) in the mean spike number in the four W-P transition pathways from **b**. The pathway slopes are reported next to every population panel. Top two values are for “Solid” lines (red square for red, blue square for blue), and the bottom two values are for “Dash” lines. (d) A map illustrates the four analysed networks; black dot: initial connectivity (W1P1); orange dot: initial weights with doubled density (W1P2); blue dot: doubled weights with initial density (W2P1); pink dot: doubled weights and density (W2P2). (e) Line plots (top, L4E; bottom, L4I) show a temporal evolution of the mean spike number normalised by the population size (<#Spikes>/N; mean in solid ± SEM in shade). The activity is followed in time from the peak in thalamic population spiking (i.e., input onset, set at 0 ms) to 60 ms post-onset (bin size: 3 ms). Colour codes match the legend in **d.** (f) Linearity of the interaction between the effects of W and P from **e**, for L4E (top) and L4I (bottom). The difference between spiking in W2P2 (pink in c) was divided by the summed effect of W1P2 (orange in c) and W2P1 (blue in c) as: W2P2 / (W2P1+W1P2). Grey area (supra): supralinearity; white area (sub): sublinearity. (g) Distributions of structural embedding scores for L4E (left) and L4I (right) neurons. For each neuron, the score (horizontal) was calculated as the sum over its K presynaptic partners of synaptic weight (W_syn_) multiplied by maximal conductance (g_syn_). Vertical axes, percentage of neurons. Distributions correspond to networks from panel d (see legend in L4I panel).

### 4.2. Weight-Dependent Effects of Density Scaling Drive Bidirectional Activity Control

To dissect the interplay between W and P, we evaluated the responses of L4E and L4I populations along “W–P pathways” where one parameter was held fixed and the other was incrementally upscaled (Figure 2b). Doubling W at either baseline (P1) or increased P (P2) boosted L4E spikes by 300% and 250% (slopes: 2.56, 1.94; Figure 2c), and L4I by 200% and 100% (slopes: 1.99, 0.76), relative to W1P1. The effect of W on L4E was independent of P (red-solid vs red-dashed slopes: p=0.292, linear regression with interaction term), while L4I activity could be enhanced more under P1 and significantly less under P2 (p=0.017). Similar to analysing the effects of upscaling W at fixed P, increasing P at W1 had almost no impact (slopes: L4E −0.04, L4I −0.08), but at W2, it *reduced* L4E and L4I spikes (slopes: −0.39, −1.10), thus exhibiting bidirectional control over spiking. This effect was stronger in L4I (blue-solid vs blue-dashed: p=0.339, L4E; p<0.005, L4I). These results define a monotonic influence for W, but a nuanced, W-dependent modulation of activity by P.

### 4.3. Sublinear Weight-Density Interactions Arise from Inhibitory Stabilisation

To clarify the interaction between W and P over time, we tracked spiking in four network states: baseline (W1P1), high W (W2P1), high P (W1P2), and combined W-P upscaling (W2P2) (Figure 2d). Baseline networks showed transient response peaks; enhanced W prolonged and elevated responses, while increased P selectively suppressed early activity (Figure 2e). Combined upscaling of W and P (W2P2) produced lower activity than W2P1 (Table 1), suggesting a ceiling effect. Quantifying this with the linearity score L = W2P2 / (W2P1+W1P2), we observed consistent sublinear integration (L<1) for both L4E and L4I (Figure 2f). Sublinear integration coincided with increasingly negative structural embedding scores (calculated as neuron-specific indegree weighted by the synapse-specific weight and maximal conductance, see Supplementary Materials) as network density and synaptic strength increased (Figure 2g, p<0.001, one-way ANOVA), suggesting inhibitory stabilisation limits joint W–P effects.

**Table 1.** Mean L4 spiking activity from early (0-20 ms) and late (40-60 ms) response periods relative to the onset of thalamic activity, shown in Figure 2e. Activity, defined as the mean number of spikes per neuron (<#Spikes>/N), is reported as mean ± standard deviation (SD). All bold values have p < 0.001 after Wilcoxon signed-rank tests vs. baseline (W1P1), Bonferroni-corrected.

| Time window /<br>*Population* | W1P1 | W1P2 | W2P1 | W2P2 |
| --- | --- | --- | --- | --- |
| 0–20 ms, *L4E* | 0.25 ± 0.07 | **0.18 ± 0.06** | **0.30 ± 0.03** | 0.25 ± 0.07 |
| 0–20 ms, *L4I* | 0.28 ± 0.08 | **0.21 ± 0.07** | 0.30 ± 0.03 | **0.20 ± 0.06** |
| 40–60 ms, *L4E* | 0.07 ± 0.10 | 0.06 ± 0.07 | **0.24 ± 0.08** | **0.18 ± 0.10** |
| 40–60 ms, *L4I* | 0.09 ± 0.11 | 0.09 ± 0.09 | **0.26 ± 0.09** | **0.14 ± 0.07** |

### 4.4. Synaptic Weight Scaling Disrupts Excitatory While Enhancing Inhibitory Cross-Laminar Spiking Rate Transmission

Analysis of spiking in L2/3, the recipient of L4 input, showed activity maps resembling L4 but with greater sensitivity to W (Figure 3a). Specifically, W–P pathway slopes were steeper in L2/3E (2.73 at P1; Figure 3b) and L2/3I (3.22) than in corresponding L4 populations. At high P, slopes remained higher in L2/3 (E: 1.91; I: 2.45). Firing rates in L2/3 increased up to 1290% for E and 500% for I under strong W scaling, exceeding rate increases in L4. Bidirectional activity modulation by P persisted, especially in high W regimes (Figure 3b), while joint scaling of W and P remained sublinear (Supplementary Figure 3a-b), and correlated with more negative embedding scores compared to W1P1 (Supplementary Figure 3c, p<0.001, ANOVA).

**Figure 3:**
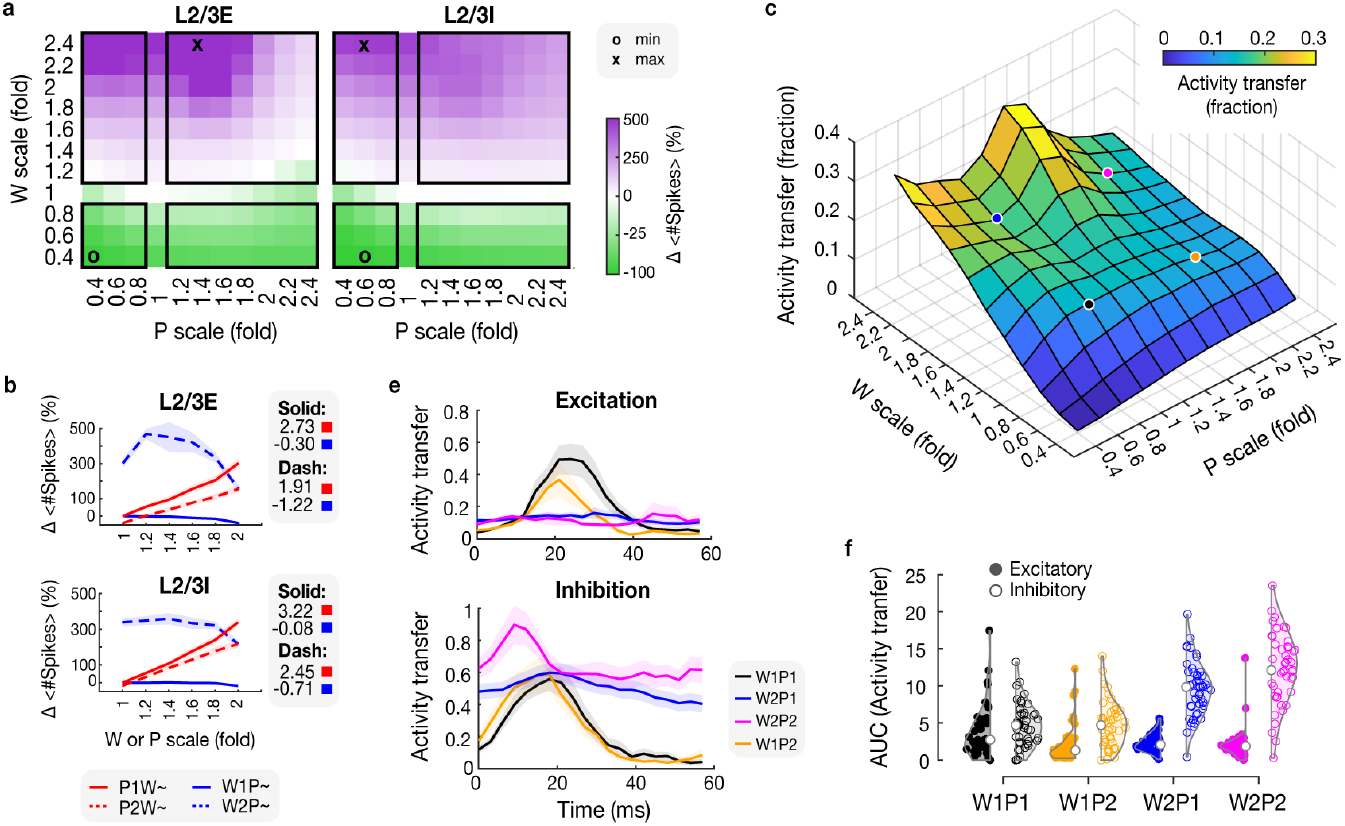
Modulation of spiking activity in the superficial layer 2/3 and the cross-laminar activity transfer changes after scaling synaptic weight and density. (a) Heatmaps for the activity of the L2/3 excitatory (E, left map) and inhibitory (I, right map) populations show the relative change (%) in the mean spike number (#) with respect to W1P1 (colour scale on the right, shared for both maps). Synaptic weights (W, vertical axis) and density (P, horizontal axis) were scaled symmetrically (scaling factors: 0.4-2.4, increments of 0.2). Symbols on maps point to conditions with the maximal positive change (x) and the maximal negative change (o). (b) Line plots (top, L2/3E; bottom, L2/3I) show a relative change (%) in the mean spike number in the four W-P transition pathways (see legend below, also Figure 2b). The pathway slopes are reported next to every population panel. Top two values are for “Solid” lines (red square for red, blue square for blue), and the bottom two values are for “Dash” lines. (c) A fraction of transferred activity (calculated as the sum of spikes in L2/3 divided by the sum of spikes in L4, colorbar in top right and height axis) from L4 to L2/3, at different weight (W, depth axis) and density (P, width axis) scales. (d) Fraction of the excitatory (top) and inhibitory (bottom) activity transfer (vertical axes) in time since a thalamic event (horizontal axes), calculated as the sum of L2/3 spikes per time bin (3 ms) divided by the sum of L4 spikes per bin, in the four analysed networks: black, W1P1; blue, W2P1; orange, W1P2; pink, W2P2 (see legend). (e) Change in total activity transfer (vertical) quantified as the area under the curve (AUC) for excitatory (left violins and filled circles) and inhibitory (right violins and open circles) activity transfer fractions from **d**. The horizontal axis shows the different connectivity scaling conditions.

We quantified cross-laminar “transmission gain” as the mean L2/3 firing divided by the L4 mean. The full W-P scaling landscape showed attenuation (gain<1), with similar values in the main conditions (W1P1: 15±10%, W2P1: 18±7%, W1P2: 15±16%, W2P2: 16±7%; W1P2 vs W2P1 differed, p=0.018; Figure 3c). At baseline, event-locked peaks in transmission reflected precisely timed propagation of excitation and inhibition (Figure 3d). Increasing P selectively reduced excitatory gain (area under the curve (AUC): 3.84±3.60, W1P1; 2.32±2.62, W1P2; p=0.006, one-way ANOVA; Figure 3e). Conversely, increasing W had broad, neuron-specific effects: the excitatory peak was eliminated and the overall gain decreased (AUC: 2.35±1.16, W2P1), while the inhibitory peak remained and the overall gain increased (AUC: 4.68±2.95, W1P1; 9.59±3.70, W2P1; p<0.005, one-way ANOVA). Thus, W controls precision and magnitude in cross-laminar spike transmission, reducing excitatory temporal structure and enhancing inhibitory signalling.

### 4.5. Layer-Specific Neuronal Recruitment Dynamics Expose Opposing Weight-Dependence Between L2/3 and L4

We analysed how W and P scaling affected the dynamics of active neurons by employing three separate metrics: the active pool size, defined as the fraction of neurons active per stimulus, the recruitment likelihood (RL), defined as the fraction of stimuli that activate a neuron, and the firing rate changes of active (recruited) neurons.

At baseline (W1P1), approximately half of L4 neurons were active per stimulus (E: 48.4%, I: 49.2%), whereas L2/3 pools were substantially smaller (E: 13.4%, I: 33.2%). Connectivity scaling produced distinct active pool dynamics across layers (Figure 4a). Under high W and low P, L2/3 demonstrated large expansion capacity (maxima: +230% E, +89% I), exceeding L4 (maxima: +47% E, +39% I). Pool size was minimised under opposite W–P regimes: in L4 at high W and high P, and in L2/3 at low W and low P.

**Figure 4:**
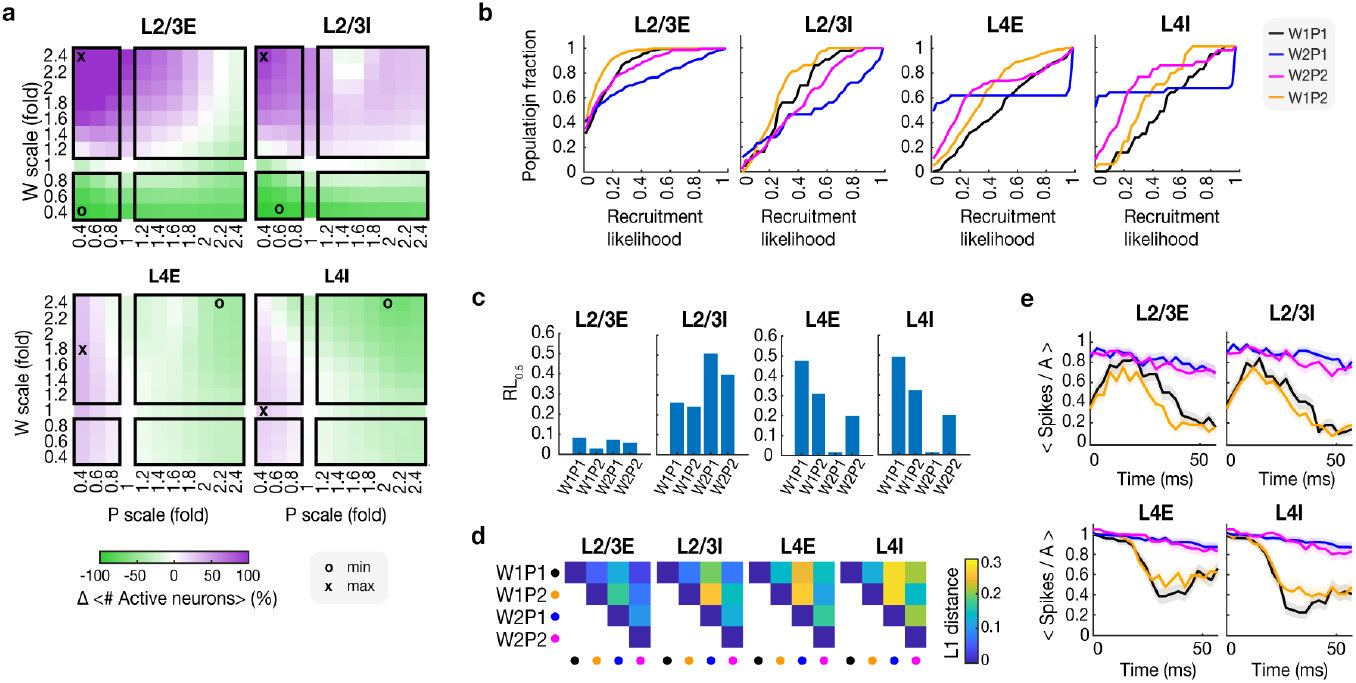
Changes in recruitment likelihoods and activity rates of single neurons after synaptic weight and density scaling. (a) Relative change (%) in the average number of active neurons per stimulus window (Methods) with respect to W1P1 in L2/3E (top left heatmap), L2/3I (top right), L4E (bottom left), L4I (bottom right). The colour scale below L4E is shared for all maps. Scaling of W, vertical axis; scaling of P, horizontal axis. x, maximal positive change; o, maximal negative change. (b) Cumulative distributions showing the fraction of neurons (vertical) with a given recruitment likelihood (horizontal), defined as the fraction of stimuli that elicited activity in each neuron. Panels correspond to populations (in titles): first, L2/3E; second, L2/3I; third, L4E; fourth, L4I. Distribution colours indicate four networks (see legend next to L4I). (c) Median recruitment likelihood (RL_0.5_, vertical), defined as the recruitment likelihood at 50% of the population from the cumulative distributions in **b**, for the four populations (panels: first, L2/3E; second, L2/3I; third, L4E; fourth, L4I) and four analysed networks (horizontal: W1P1, W1P2, W2P1, W2P2). (d) Manhattan (L1) distances, quantifying the total difference between cumulative distributions in **b**, are shown for all network pairs. Network identities are indicated by marker colour (black: W1P1; orange: W1P2; blue: W2P1; pink: W2P2) on the horizontal and vertical axes. Panels correspond to populations (in titles): first, L2/3E; second, L2/3I; third, L4E; fourth, L4I. L_1_ distance values are represented by the colour scale (see legend next to L4I). (e) The temporal evolution of the firing rates of active (A) neurons (mean in solid ± SEM in shade) from four networks: black, W1P1; orange, W1P2; blue, W2P1; pink, W2P2 (see legend next to L4I **b**). The activity is measured from the input pattern onset (0 ms) to 60 ms post-onset (bin size, 3 ms). Population names are above the panels.

Recruitment likelihood distributions corroborated these layer-specific patterns (Figure 4b). At baseline, L4 exhibited broad, approximately uniform RL distributions, with median recruitment (RL_0.5_) values near 0.5 (L4E 0.48; L4I 0.49; Figure 4c), whereas L2/3 distributions were narrower with lower RL_0.5_ (L2/3E 0.08; L2/3I 0.26). Increasing P shifted all distributions toward lower RL, reducing RL_0.5_ to 0.31 (L4E), 0.33 (L4I), 0.03 (L2/3E), and 0.24 (L2/3I), consistent with smaller pool sizes. Weight potentiation (W2P1) exerted the largest reshaping effect on recruitment distributions, as quantified by Manhattan (L1) distances comparing cumulative distributions (Figure 4d, Supplementary Materials). In L2/3, W2P1 increased RL_0.5_ for inhibitory neurons (to 0.51) and broadened the upper tail of the excitatory distribution despite minimal change in its RL_0.5_, indicating selective amplification of a subpopulation of E neurons. In L4, W2P1 sharply reduced RL_0.5_ (E 0.02; I 0.01) and yielded a bimodal pattern in which most neurons were rarely active but a minority responded reliably to nearly every stimulus.

Joint upscaling (W2P2) partially restored recruitment profiles toward baseline. Distributions shifted back with intermediate RL_0.5_ values (L4E 0.20; L4I 0.20; L2/3E 0.06; L2/3I 0.40; Figure 4c) and reduced L1 distances compared to W2P1 (Figure 4d). Finally, among recruited neurons, mean firing rates and response duration increased robustly with W across all populations, whereas P had comparatively minor effects (Figure 4e; W2P1 and W2P2 > W1P1 and W1P2, p<0.001, Kruskal–Wallis; Table 2). These results demonstrate that P primarily regulates recruitment (active pool size, RL), while W modulates the activity amplitude and duration of recruited neurons. Moreover, recruitment dynamics is layer-specific (expansion in L2/3 versus contraction in L4), but firig rate amplification among active neurons is not.

**Table 2.** Mean L4 and L2/3 spiking activity of recruited neurons, separated in early (0-20 ms) and late (40-60 ms) response periods relative to the onset of thalamic activity, shown in Figure 4e. Activity, defined as the mean number of spikes per active (A) neuron (<#spikes>/A), is reported as mean ± standard deviation (SD). Bold values show p < 0.001, statistical comparisons between groups, the Kruskal–Wallis test with Bonferroni correction.

| Time window /<br>*Population* | W1P1 | W1P2 | W2P1 | W2P2 |
| --- | --- | --- | --- | --- |
| 0–20 ms, *L2/3E* | 0.67 ± 0.47 | 0.57 ± 0.50 | **0.89 ± 0.31** | **0.89 ± 0.31** |
| 0–20 ms, *L2/3I* | 0.65 ± 0.48 | 0.58 ± 0.49 | **0.93 ± 0.26** | **0.91 ± 0.30** |
| 0–20 ms, *L4E* | 0.95 ± 0.22 | 0.98 ± 0.15 | **0.99 ± 0.15** | **1.02 ± 0.13** |
| 0–20 ms, *L4I* | 0.93 ± 0.27 | 0.94 ± 0.25 | **0.99 ± 0.15** | **1.01 ± 0.20** |
| 40–60 ms, *L2/3E* | 0.22 ± 0.42 | 0.17 ± 0.38 | **0.75 ± 0.43** | **0.71 ± 0.46** |
| 40–60 ms, *L2/3I* | 0.14 ± 0.34 | 0.14 ± 0.35 | **0.82 ± 0.39** | **0.74 ± 0.44** |
| 40–60 ms, *L4E* | 0.56 ± 0.50 | 0.58 ± 0.49 | **0.89 ± 0.31** | **0.85 ± 0.37** |
| 40–60 ms, *L4I* | 0.40 ± 0.49 | 0.43 ± 0.50 | **0.89 ± 0.32** | **0.83 ± 0.42** |

**Table 3.** Connectivity parameters. Synaptic weights (W) and connectivity probabilities (P) are specific per projection. A projection type is defined as a collection of synapses with identical pre- and post-population pairs (e.g., L4E → L4I). W and P values are taken from **(Markram et al. 2015)**.

| Pre-post layer pairs | Pre-post neuron pairs | W (unitless) | P (%) |
| --- | --- | --- | --- |
| *L2/3* → *L2/3* | *E* → *E* | 2.8 | 5.6 |
|  | *E* → *I* | 8.1 | 5.1 |
|  | *I* → *E* | 17 | 5.9 |
|  | *I* → *I* | 15 | 5.1 |
| *L4* → *L2/3* | *E* → *E* | 2.4 | 5.8 |
|  | *E* → *I* | 5.7 | 0.33 |
|  | *I* → *E* | 14 | 1.7 |
| *L4* → *L4* | *E* → *E* | 3.3 | 7.6 |
|  | *E* → *I* | 7.9 | 4.2 |
|  | *I* → *E* | 16 | 6.3 |
|  | *I* → *I* | 14 | 6.2 |
| *VPM (pattern-generating)* → *L4* | *E* → *E* | 4.55 | 2.79 |
|  | *E* → *I* | 5.85 | 3.25 |
| *VPM (back-ground)* → *L4* | *E* → *E* | 2.45 | 1.5 |
|  | *E* → *I* | 3.15 | 1.75 |

### 4.6. Connectivity Scaling Reveals Cell-Specific E/I Balance Regimes Underlying Neuronal Functional Classes

To understand the basis of altered neuronal recruitment into active pools following connectivity scaling, neurons were classified across the four network states (W1P1, W1P2, W2P1, W2P2) by responsiveness to thalamic drive (Figure 5a), yielding three functional classes: scaling-invariant (active in all states), scaling-variant (active in a subset), and silent (inactive in all states). All populations contained invariant and variant neurons, with silent cells emerging in L2/3 (Figure 5b; L2/3E: 45.9% invariant, 49.6% variant, 4.5% silent; L2/3I: 83.7% invariant, 16.3% variant; L4E: 55.9% invariant, 44.1% variant; L4I: 63.6% invariant, 36.4% variant). This classification suggests functional heterogeneity that is universal across all populations, with two complementary roles for neuronal subpopulations: invariant neurons maintain baseline responsiveness across W–P regimes, whereas variant neurons support adaptability by altering recruitment with network state.

**Figure 5:**
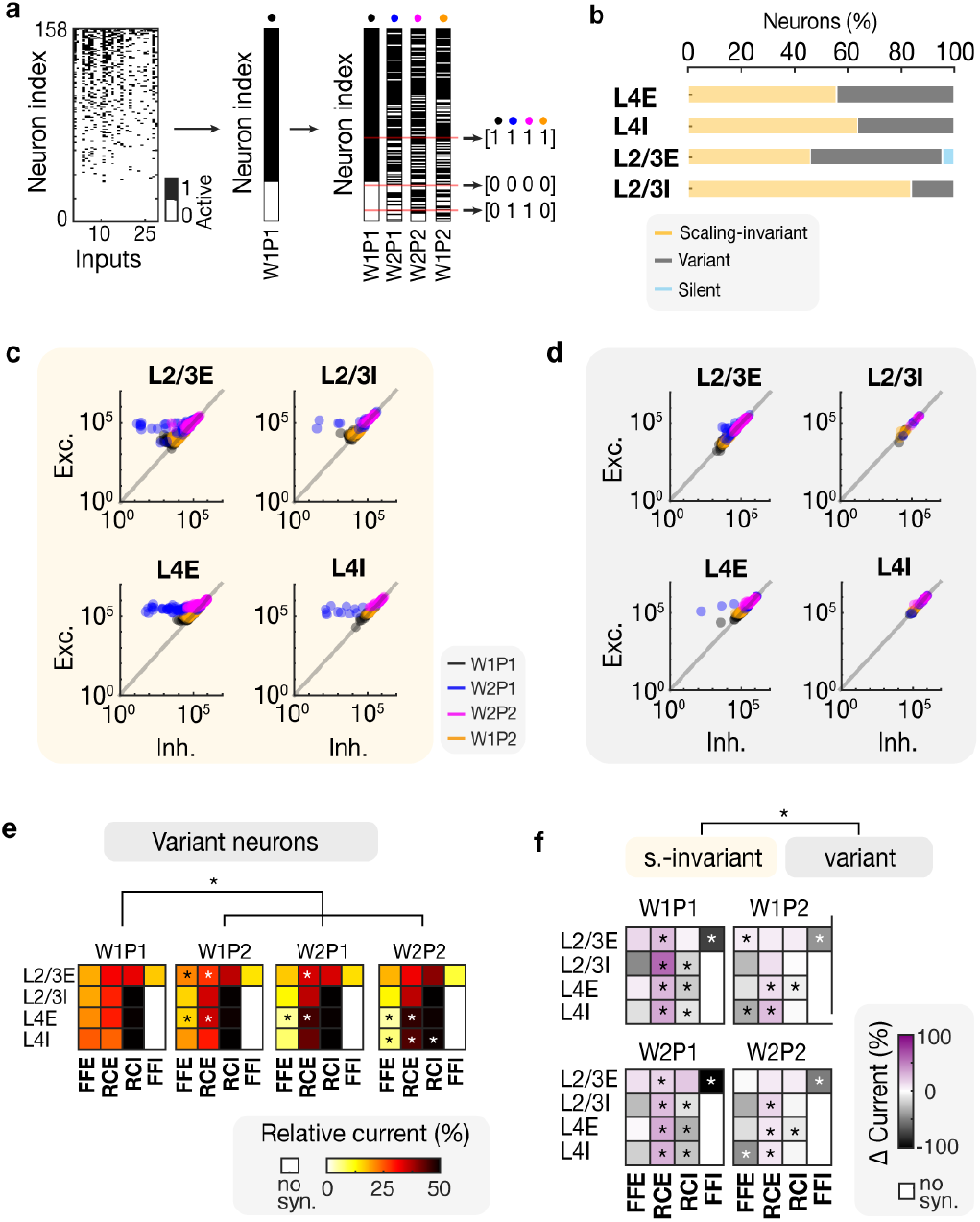
The impact of increased weight and connection density on the excitation-inhibition balance in distinct neuronal subpopulations. (a) Illustrated example of the classification protocol behind data in **b**. Left: inputs (horizontal axis, Methods) presented to the network triggered activity (black points) in neurons (vertical axis). Middle: activity from the population (left) represented as a binary vector: neurons responsive to ≥ 1 thalamic event get assigned a 1 (black) and a 0 otherwise (white). Right: four binary vectors represent the same population (left) across four different network states (see coloured dots on top or labels below each vector): black, W1P1; orange, W1P2; blue, W2P1; pink, W2P2. Numbers on the right: classification protocol with three categories (examples shaded with red lines): (1) neurons responsive in all four conditions (code: 1111, label: scaling-invariant), (b) Functional categorisation of neurons based on their responsiveness across four network states (in labels, bottom right; Methods) produced three classes: invariant (responded in all network states, orange); variant (responded in some network states, grey); silent (unresponsive in all states, blue). Stacked bars show the percentage (%, vertical) of each class in the four populations (L4E, L4I, L23E, L23I). (c) For each invariant neuron (orange in b), the sum of all excitatory (vertical axis) and all inhibitory currents (horizontal axis) within the stimulus window (Methods) is shown. Dots are neurons, colours are the four conditions (black, W1P1; blue, W2P1; pink, W2P2; orange, W1P2), and panels are populations (see titles). (d) Same as in **c**, but for variant neurons (grey fractions in **b**). (e) Relative contributions of the four presynaptic current types to variant neurons (horizontal axis: FFE, feedforward excitation; RCE, recurrent excitation; RCI, recurrent inhibition; FFI, feedforward inhibition) are shown for each population (vertical axis: L4E, L4I, L23E, L23I). Heatmaps are arranged by condition (titles): first, W1P1; second, W1P2; third, W2P1; fourth, W2P2. Relative contribution represents the percentage of total current received by each neuron and is indicated by the colorbar below. L4E, L4I, and L2/3I lack FFI inputs (colorbar: “no syn.”). Asterisks denote significant differences per current type compared to W1P1, based on Wilcoxon signed-rank tests with Bonferroni correction (^*^, p < 0.05). (f) Differences in relative current subtypes (horizontal axis: FFE, feedforward excitation; RCE, recurrent excitation; RCI, recurrent inhibition; FFI, feedforward inhibition) between input-invariant and variant neurons across populations (vertical axis: L4E, L4I, L23E, L23I) and conditions: top left, W1P1; top right, W1P2; bottom left, W1P2; bottom right, W2P2. Colour scale (below) applies to all maps. Asterisks denote significant differences per current type compared between invariant and variant neurons, based on Wilcoxon signed-rank tests with Bonferroni correction (^*^, p < 0.05).

E/I balance differed systematically between classes and scaled with connectivity. Using neuron-specific log(E/I), we quantified the fraction of near-balanced neurons (|log(E/I)| < 0.5) per population and network state. The balanced fraction was lower among invariant neurons at baseline (W1P1), reflecting heterogeneity in E/I balance (L2/3E 0.57; L2/3I 0.67; L4E 0.57; L4I 0.81; Figure 5c), but was uniformly high among variant neurons (>0.9 across all populations; Figure 5d), indicating tightly regulated balance. Connectivity scaling primarily affected the E/I balance of scaling-invariant neurons, while variant neurons remained largely stable. Specifically, in invariant neurons, upscaling P increased the balanced fraction (L2/3E 0.92; L2/3I 0.89; L4E 0.86; L4I 1.00), while upscaling W decreased it as neurons shifted towards excitation-dominated regimes, particularly in L4 (Figure 5c; (L2/3E 0.64; L2/3I 0.69; L4E 0.34; L4I 0.33). Joint upscaling (W2P2) partially restored the balance fraction in the invariant class toward baseline (L2/3E 0.89; L2/3I 0.94; L4E 0.61; L4I 0.71). Variant neurons remained tightly balanced under all conditions (>0.9), suggesting a class-specific, stable E/I operating point.

Decomposing presynaptic inputs into feedforward excitation and inhibition (FFE, FFI; synapses from the preceding layer) and recurrent excitation and inhibition (RCE, RCI; synapses from the same layer) revealed the circuit basis for these shifts. Variant neurons sustained balance through consistently strong inhibition across states, with approximately half of the total presynaptic input inhibitory at baseline (Figure 5e). Inhibitory drive was dominated by RCI in L2/3I and in L4 neurons, and by combined RCI and FFI in L2/3E. Weight upscaling redistributed excitatory sources in all populations from a nearly uniform composition at baseline (FFE ≈ RCE) toward stronger RCE and weaker FFI (Figure 5e; p < 0.05 for currents marked with an asterisk, Wilcoxon signed-rank test), yet overall inhibition remained robust in variants, preserving balance despite changes in excitatory pathways.

Comparing classes directly revealed that invariant neurons received weaker inhibition (RCI and FFI) and stronger RCE than variants across all states, with differences most pronounced in W1P1 and W2P1 (Figure 5f; p < 0.05 for currents marked with an asterisk, Wilcoxon signed-rank test). Across the full W-P plane, invariant neurons remained under markedly weaker inhibitory control than variants (Supplementary Figure 4a-b), explaining their persistent recruitment and heightened sensitivity to weight scaling.

Altogether, these analyses indicate that presynaptic current composition sets the functional class: scaling-invariant neurons are characterised by high recurrent excitation with reduced recurrent and feedforward inhibition, leading to excitation-dominated regimes that ensure consistent recruitment; scaling-variant neurons maintain tight E/I balance via strong recurrent inhibition that covaries with excitation, resulting in a probabilistic, state-dependent recruitment. Weight scaling alone pushes invariant neurons deeper into excitation dominance, whereas density scaling recruits proportional inhibition to restore balance. Consequently, W-P interactions define cell-specific E/I set points that predict and govern single-cell recruitment patterns across connectivity regimes.

## 4. Discussion

Neuronal intrinsic properties interact with two fundamental features of network connectivity — synaptic weight (W), which sets pairwise connection strength, and connection probability (P), which governs network sparsity — to generate complex population dynamics. This work advances the understanding of how W and P jointly shape activity regimes in biological, nonlinear circuits. Our data show that synaptic weight dominates in setting the overall network spiking, while connection sparsity modulates spiking dynamics bidirectionally, in a weight-dependent manner. Scaling these structural features reveals distinct computational modes and functionally divergent neuronal subtypes.

### 4.1. Synaptic Weight as the Primary Driver of Network Dynamics

Upscaling W strongly amplifies firing rates, prolongs responses, and shifts cross-laminar activity gain, identifying synaptic efficacy as the main determinant of network activity and its dynamic range. This aligns with homeostatic synaptic scaling that regulates postsynaptic efficacy to stabilise firing (Turrigiano 2012) and with Hebbian and STDP-like processes that modulate W and explain rate variability in cortex (Celikel et al. 2004; Celikel and Sakmann 2007). Theoretical studies predict higher activity rates for networks with larger mean (Iyer et al. 2013) and outline regimes where sparse connectivity with strong synapses reproduces cortical-like activity (Brunel 2000; van Vreeswijk and Sompolinsky 1996; 1998). Consistent with this view, in our network, stronger W paired with sparser P extend the network’s activity range, while dense networks with weak synapses approach quiescence, matching classical balanced-network predictions.

### 4.2. Connection Sparsity Provides Bidirectional Control

Unlike the largely monotonic effect of W, P exerts nuanced, weight-dependent control over population output. In standard connectivity scaling approaches (Brunel 2000; van Vreeswijk and Sompolinsky 1998), P is often absorbed into W to preserve network dynamics, such as first- and higher-order statistics (van Albada et al. 2015), which masks its functional role. Decoupling W and P in our network demonstrates that at low W, increasing P produces minimal change in spiking, whereas at high W, the same P increase suppresses activity. This bidirectionality situates P as a structural gain control that can either preserve or attenuate output depending on the synaptic-weight regime, suggesting a mechanism by which connectopathies — disorders of network wiring (van den Heuvel and Sporns 2019) — could differentially modulate circuit function contingent on concurrent plasticity of W.

### 4.3. Nonlinear Weight–Sparsity Interactions and Inhibitory Stabilisation

Joint upscaling of W and P consistently yielded sublinear integration: the response under combined scaling was less than the sum of responses after individual W and P scalings. This pattern coincides with inhibitory stabilisation, thus suggesting that in our network, denser and stronger synapses preferentially recruit feedback inhibition. Inhibitory stabilisation was previously suggested as a control mechanism for plasticity-driven changes in weights (Naumann and Sprekeler 2020). While Naumann and Sprekeler (2020) define *how* presynaptic inhibitory activity could downscale excitatory synaptic efficacy, we map the weight-density combinations *where* such inhibition is recruited. Although P is unlikely to change on fast timescales required for immediate stabilisation, episodes of elevated excitation under high W could trigger slower structural adjustments that increase P, moving the circuit toward regimes where inhibitory stabilisation attenuates subsequent hyperexcitability. This two-timescale mechanism — rapid weight-driven excitation followed by slower density-mediated stabilisation — provides a concrete interaction between synaptic and structural homeostasis.

### 4.4. E/I Regimes Underlying Single Neuron Responsiveness

Our results identify a division between scaling-invariant neurons, which are persistently recruited across all network states, and scaling-variant neurons, which exhibit state-dependent recruitment with tightly balanced E/I currents. This mirrors recent data from both modelling and transcriptomic studies showing that subsets of cortical neurons sustain consistent firing across changing conditions due to reduced inhibitory tone, while others maintain tight E/I balance and display probabilistic activity, contributing to population code heterogeneity (Denève and Machens 2016; Sukenik et al. 2021; Barrett et al. 2016). Studies of efficient coding and inhibitory control corroborate that recurrent inhibition modulates variability and defines recruitment thresholds (Barrett et al. 2016; Festa et al. 2021), supporting our conclusion that variation in presynaptic current ratios determines whether a cell acts as a stable responder or a flexible participant in dynamic ensembles.

### 4.5. Implications for cortical computation and plasticity

Together, these findings show that the interplay between W, P, and inhibition governs not only overall network activity but also functional differentiation among neurons across layers. By linking structural parameters to dynamic specialisation, our model offers a framework for understanding how cortical circuits balance stability with adaptability. Neurons with weaker recurrent inhibition may constitute initial loci of homeostatic or Hebbian plasticity, while those under strong inhibition provide a stable scaffold that maintains population balance. This division could allow learning to proceed locally without global destabilisation. Moreover, the differential sensitivity of layers and neuronal classes to W and P suggests that cortical circuits exploit structure to tune responsiveness and selectivity. In summary, W and P jointly define a continuum between stability and adaptability, providing a structural substrate through which the cortex can flexibly regulate computation, plasticity, and resilience to perturbation.

## 5. Acknowledgements

This work was supported by a grant from the Netherlands Organisation for Scientific Research (NWO Vidi grant VI.Vidi.213. 137) to F.Z. and an internal grant from the Donders Institute for Brain, Cognition and Behaviour, Radboud University, to T.C.

## 9. Supplementary Methods

### 9.1. Cortical Neurons

We used Hodgkin-Huxley models for regular spiking and fast-spiking rat somatosensory neurons from (Pospischil et al. 2008) to model excitatory and inhibitory cortical neurons, respectively (Methods 3.2.1). Membrane potential was given by:

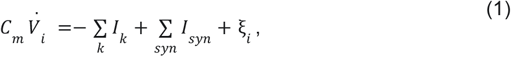

where

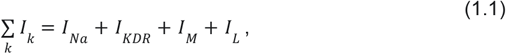

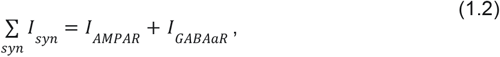

and

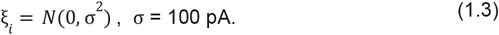

Ionic channel currents followed the generic form:

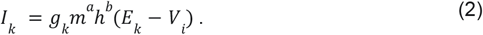

For the sodium (Na) current:

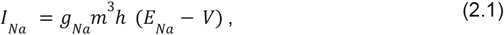

where

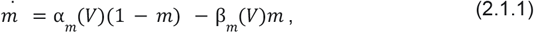

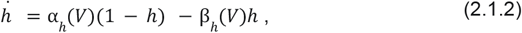

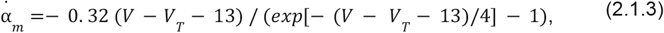

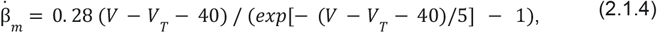

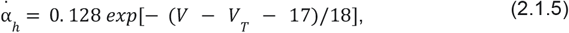

and

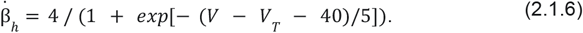

For the delayed-rectified potassium (KDR) current:

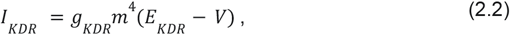

where

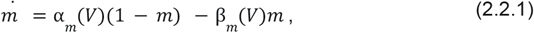

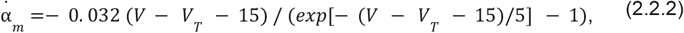

and

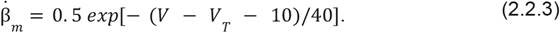

For the slow non-inactivation (M) current:

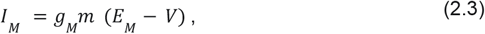

where

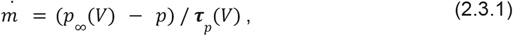

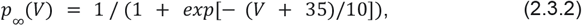

and

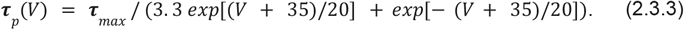

For the leak (L) current:

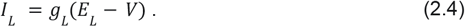

Neuron-type-specific parameter values for Eqs. 2.1 — 2.4 are reported in Supplementary Table 1.

### 9.2. Thalamic Neurons

We used Hodgkin-Huxley models for thalamocortical relay (TCR) cells from (Ching et al. 2010) to model excitatory thalamic neurons (Methods 3.2.2). Membrane potential was given by:

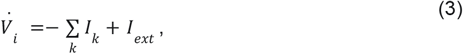

where

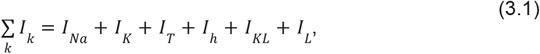

and

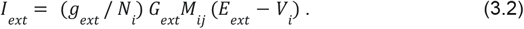

For the sodium (Na) current:

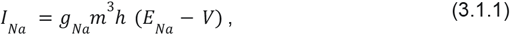

where

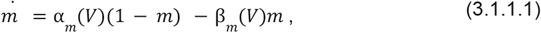

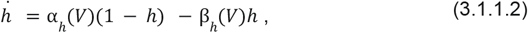

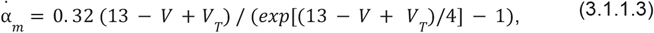

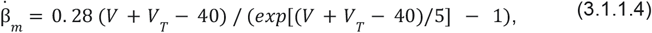

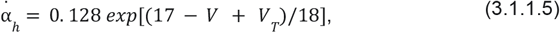

and

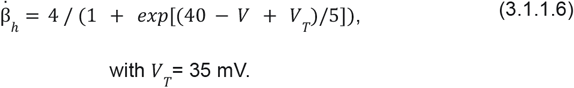

For the potassium (K) current:

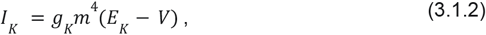

where

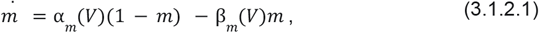

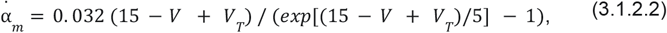

and

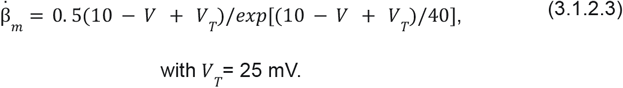

For the T-type calcium (T) current:

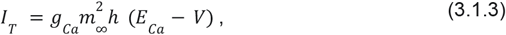

where

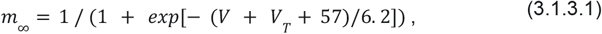

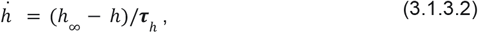

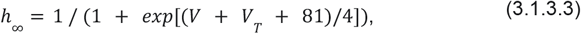

and

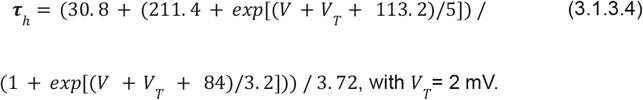

For the hyperpolarisation-activated (h) current:

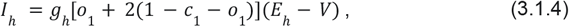

where

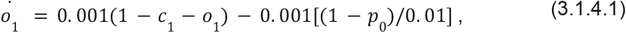

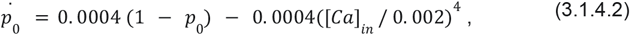

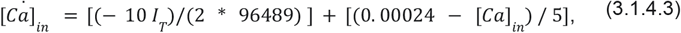

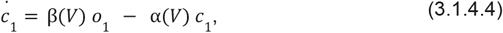

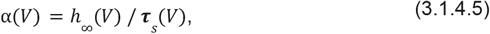

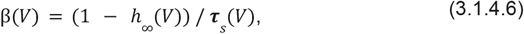

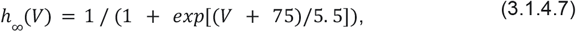

and

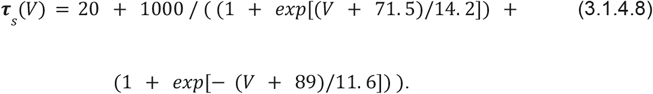

For the potassium (KL) and general leak (L) currents:

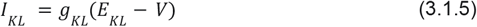

and

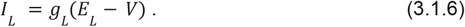

Neuron-type-specific parameter values for Eqs. 3.1.1 — 3.1.6 are reported in Supplementary Table 2.

TCR activity was triggered by external current (Methods 3.1.2):

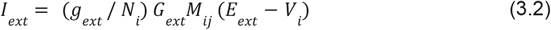

where *g*_*ext*_ was the maximal input conductance, *N*_*i*_ the number of postsynaptic neurons, *G*_*ext*_ was a binary matrix of size *N*_*i*_ × *T*, where *N*_*i*_ were rows and *T* (number of simulation timesteps) columns, with ones occurring randomly across columns based on the Poisson distribution with rate *r* (Supplementary Table 2). The binary matrix *G*_*ext*_ was convolved with an EPSP kernel *x*:

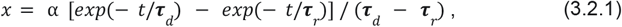

Where α was the EPSP amplitude, and ***τ***_*r*_ and ***τ***_*d*_ were the EPSP rise and decay times, respectively. The resulting input signals were neuron-specific, sharing more or less temporal correlation, depending on the thalamic subpopulation (more between pattern-generating, and less between background neurons; see Methods 3.1.2). In Eq. 3.2, *M*_*ij*_ was a binary vector with *N*_*i*_ rows and *N*_*i*_*P*_*ext*_ randomly distributed positive elements, controlling which neurons received EPSP input signals from *G*_*ext*_.

### 9.3. Synapse Model

We used the conductance-based synaptic model explained in Methods (3.3.1):

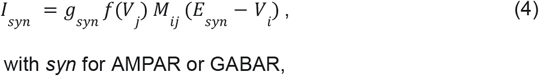

where

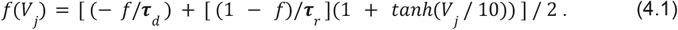

Neuron-type-specific parameter values for Eq. 4 are reported in Supplementary Table 3.

## 10. Supplementary Tables

**Supplementary Table 1.**
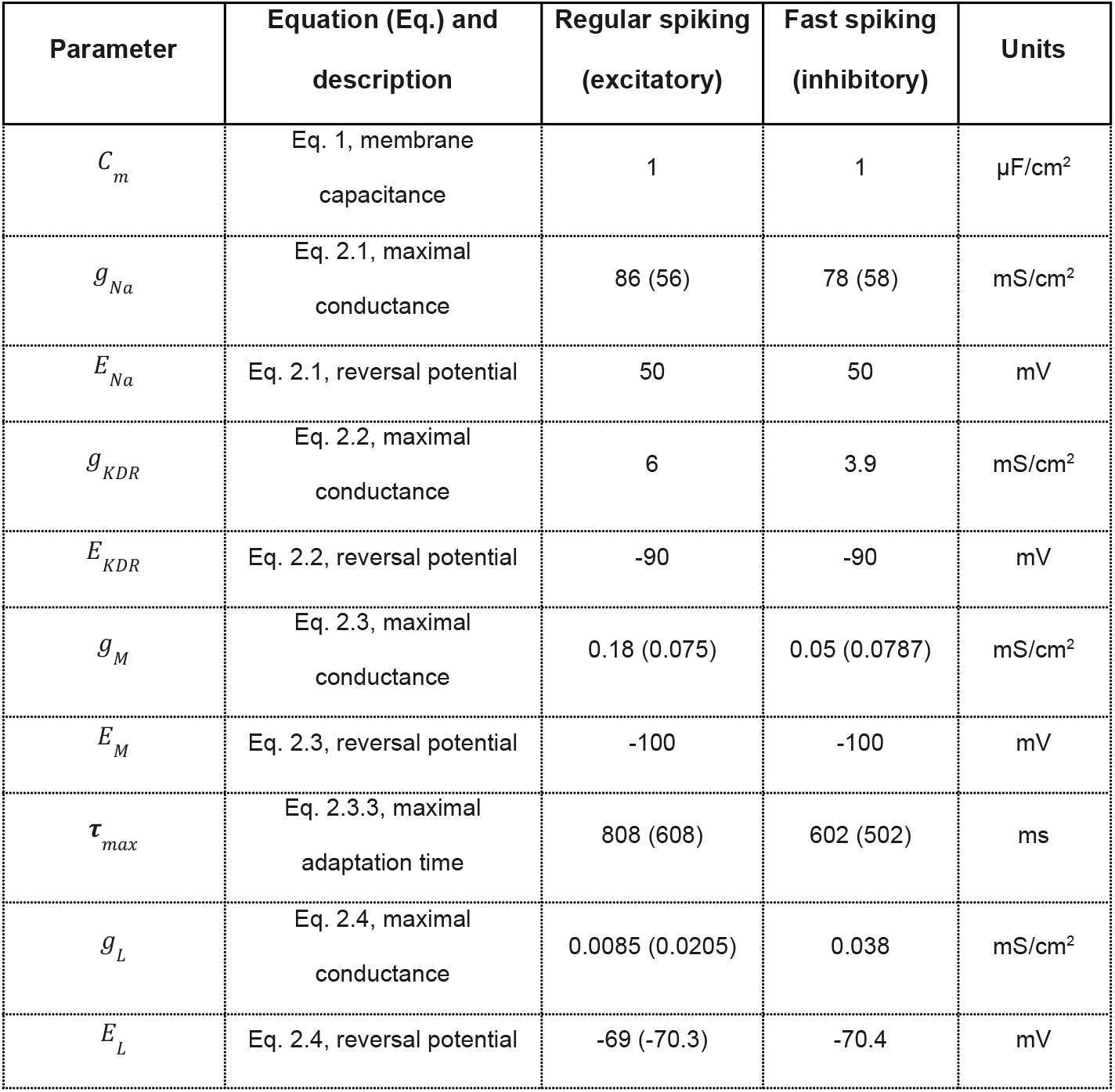
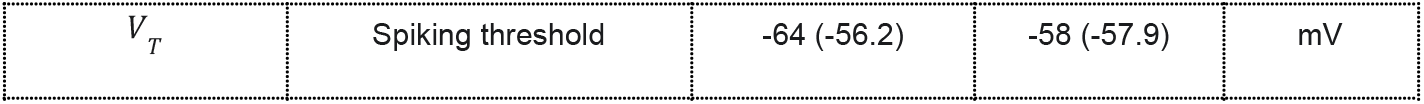
Parameter values for cortical neuron models. Values come from (Pospischil et al. 2008). If two values are reported, these were adjusted to match empirical activity rates from Supplementary Figure 1a: the adjusted value is reported outside brackets, and the original value is inside brackets.

**Supplementary Table 2.** Parameter values for thalamic (TCR) neuron models. Values come from (Ching et al. 2010). If two values are reported, these were adjusted to get the desired TCR dynamics. The adjusted value is reported outside brackets, and the original value is inside brackets.

| Parameter | Equation (Eq.) and description | Pattern-generating TCR | Background TCR | Units |
| --- | --- | --- | --- | --- |
| $g_{Na}$ | Eq. 3.1.1, maximal conductance | 90 | 90 | mS/cm<sup>2</sup> |
| $E_{Na}$ | Eq. 3.1.1, reversal potential | 50 | 50 | mV |
| $g_K$ | Eq. 3.1.2, maximal conductance | 10 | 10 | mS/cm<sup>2</sup> |
| $E_K$ | Eq. 3.1.2, reversal potential | -100 | -100 | mV |
| $g_{Ca}$ | Eq. 3.1.3, maximal conductance | 2 | 2 | mS/cm<sup>2</sup> |
| $E_{Ca}$ | Eq. 3.1.3, reversal potential | 120 | 120 | mV |
| $g_h$ | Eq. 3.1.4, maximal conductance | 0.01 (0.001) | 0.005 (0.001) | mS/cm<sup>2</sup> |
| $E_h$ | Eq. 3.1.4, reversal potential | -40 | -40 | mV |
| $g_{KL}$ | Eq. 3.1.5, maximal conductance | 0.0172 | 0.0172 | mS/cm<sup>2</sup> |
| $E_{KL}$ | Eq. 3.1.5, reversal potential | -100 | -100 | mV |
| $g_L$ | Eq. 3.1.6, maximal conductance | 0.01 | 0.01 | mS/cm<sup>2</sup> |
| $E_L$ | Eq. 3.1.6, reversal potential | -70 | -70 | mV |
| $g_{ext}$ | Eq. 3.2, maximal conductance | 0.5 (1) | 0.75 (1) | mS/cm<sup>2</sup> |
| $N_i$ | Eq. 3.2, neuron number | 45 | 24 | - |
| $E_{ext}$ | Eq. 3.2, reversal potential | 1 | 1 | mV |
| $r$ | -, Poisson rate | 9 (12) | 5 (12) | Hz |
| $P_{ext}$ | -, Poisson input probability | 0.3 (0.5) | 0.1 (0.5) | - |
| $\alpha$ | Eq. 3.2.1, EPSP amplitude | 14 (10) | 25 (10) | - |
| $\tau_r$ | Eq. 3.2.1, rise time | 0.5 | 0.5 | ms |
| $\tau_d$ | Eq. 3.2.1, decay time | 2 | 2 | ms |

**Supplementary Table 3.** Parameter values for synaptic models.

| Parameter | Equation (Eq.) and description | AMPA (excitatory) | GABAa (inhibitory) | Units |
| --- | --- | --- | --- | --- |
| $g_{syn}$ | Eq. 4, maximal conductance | Cortico-cortical: 0.017;<br>Thalamo-cortical: 0.034 | Cortico-cortical: 0.125 | mS/cm<sup>2</sup> |
| $E_{syn}$ | Eq. 4, reversal potential | 0 | -80 | mV |
| $\tau_r$ | Eq. 4.1, rise time | 0.2 | 0.2 | ms |
| $\tau_d$ | Eq. 4.1, decay time | 2 | 10 | ms |

## 11. Supplementary Figure Captions

**Supplementary Figure 1:**
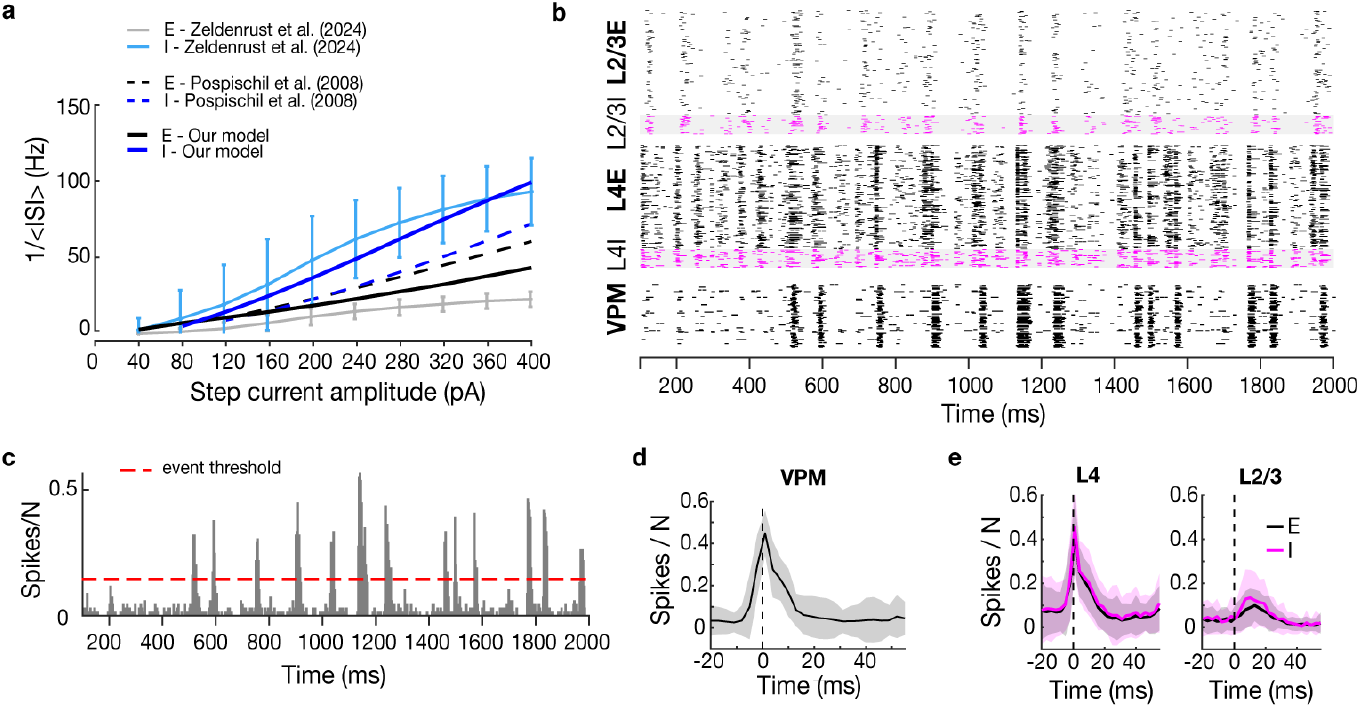
Baseline activity in a multi-layer spiking neural network model with Hodgkin-Huxley neurons representing a thalamocortical microcircuit. (a) Fitting of single-neuron dynamics for excitatory (black shades) and inhibitory (blue shades) neurons in three stages: default values from Pospischil et al. (2008), desired dynamics from Zeldenrust et al. (2024), fitted dynamics used in our model (final parameter values reported in Methods). See labels for colour codes. Vertical axis, firing rate calculated as an inverse of the mean inter-spike interval (ISI) at every current amplitude (horizontal axis). (b) Raster plot for one network realisation with initial connectivity parameters (W1P1; Methods). Vertical axis, neurons from four cortical populations (from bottom to top: L4I, L4E, L3I, L3E, and a thalamic population: VPM). Pink dots, spikes from inhibitory neurons. Black dots, spikes from excitatory neurons. The grey background behind inhibitory spikes is for better contrast. Horizontal axis, time in ms. Activity is triggered by a Poissonian process (not shown, see Methods) coupled with thalamic neurons. (c) Peristimulus time histogram (PSTH) of thalamic spiking activity (spikes per neuron, N; vertical axis) shown in panel **b** (3-ms bins; time on the horizontal axis). Randomly triggered thalamic synchronisation events served as discrete stimuli (N_stim_) for quantifying cortical responses (see panels **d**-**e**). The red line marks the detection threshold, defined as the highest peaks exceeding the mean + 1 standard deviation and separated by at least 60 ms (Methods). (d) Average thalamic PSTH (spikes per neuron, N, vertical axis; 3-ms time bins, horizontal axis) aligned to the peak synchronisation events detected as described in **c**. Activity was averaged across N_stim_ = 54 events from 5 network realisations (2s simulations each). The dashed line marks the VPM peak activity; activity is shown from 20 ms before to 40 ms after the peak. Solid line, mean; shaded area, standard deviation. (e) Average cortical PSTH (spikes per neuron, N, vertical axis; 3-ms time bins, horizontal axis) triggered by thalamic events shown in **d**. Activity is displayed by layer (L4, left; L2/3, right) and cell type: excitatory (E, black) and inhibitory (I, pink) populations. The dashed line indicates the thalamic peak activity (from **d**); activity is shown from 20 ms before to 40 ms after the peak. Solid line, mean; shaded area, standard deviation.

**Supplementary Figure 2:**
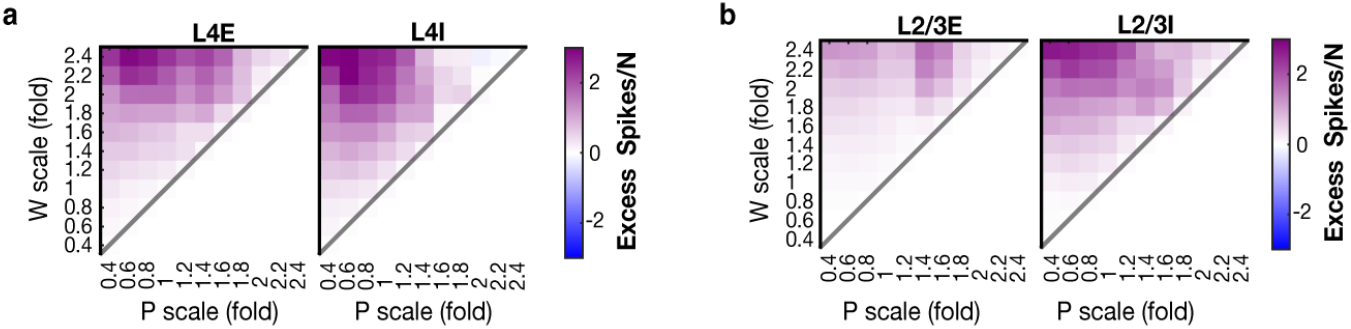
Pairwise comparison of the average evoked activity from networks with mirrored connectivity scaling: WxPy vs. WyPx. (a) Difference in evoked spikes per neuron (N) for L4E (left; see map title) and L4I (right) populations following network-wide scaling of synaptic weights (W, vertical axis) and connection probability (P, horizontal axis). Activity differences were computed between mirrored scaling conditions as WxPy − WyPx; that is, conditions below the anti-diagonal (grey) were subtracted from their counterparts above it. The colour scale (right side) represents the excess number of evoked spikes per neuron. (b) Same as in **a**, but for L2/3E (left) and L2/3I (right).

**Supplementary Figure 3:**
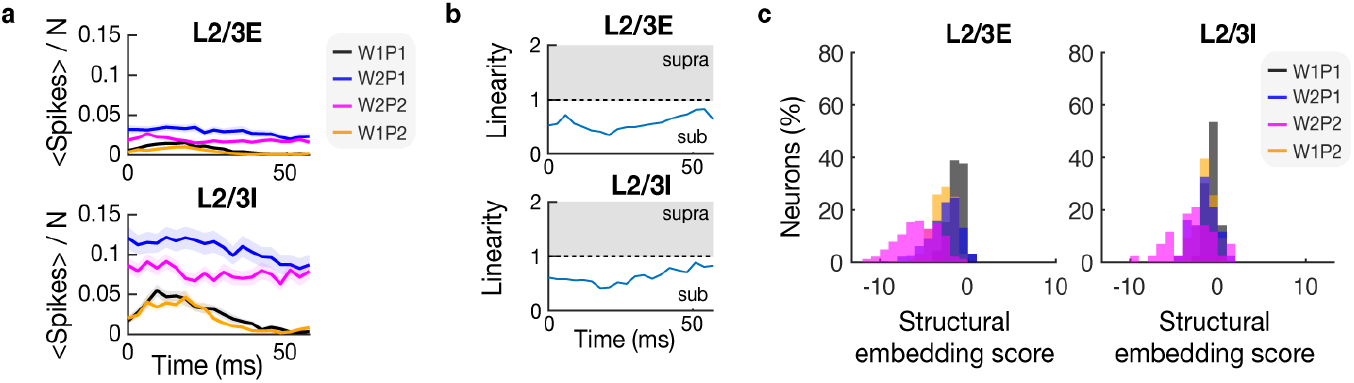
Modulation of spiking activity in the superficial layer 2/3 by changes in synaptic weights and density. (a) Line plots (top, L2/3E; bottom, L2/3I) show a temporal evolution of the mean spike number normalised by the population size (<#Spikes>/N; mean in solid ± SEM in shade). The activity is followed in time from the peak in thalamic population spiking (i.e., input onset, set at 0 ms) to 60 ms post-onset (bin size: 3 ms). Colour codes represent four network states: initial connectivity (W1P1), initial weights with doubled density (W1P2), doubled weights with initial density (W2P1), and doubled weights and density (W2P2); see legend on the right. (b) Linearity of the interaction between the effects of W and P from **a**, for L2/3E (top) and L2/3I (bottom). The difference between spiking in W2P2 (pink in **a**) was divided by the summed effect of W1P2 (orange in **a**) and W2P1 (blue in **a**) as: W2P2 / (W2P1+W1P2). Grey area (supra): supralinearity; white area (sub): sublinearity. (c) Distributions of structural embedding scores for L2/3E (left) and L2/3I (right) neurons. For each neuron, the score (horizontal) was calculated as the sum over its K presynaptic partners of synaptic weight (W_syn_) multiplied by maximal conductance (g_syn_). Vertical axes, percentage of neurons. Distributions correspond to networks from panel d (see legend in L2/3I panel).

**Supplementary Figure 4:**
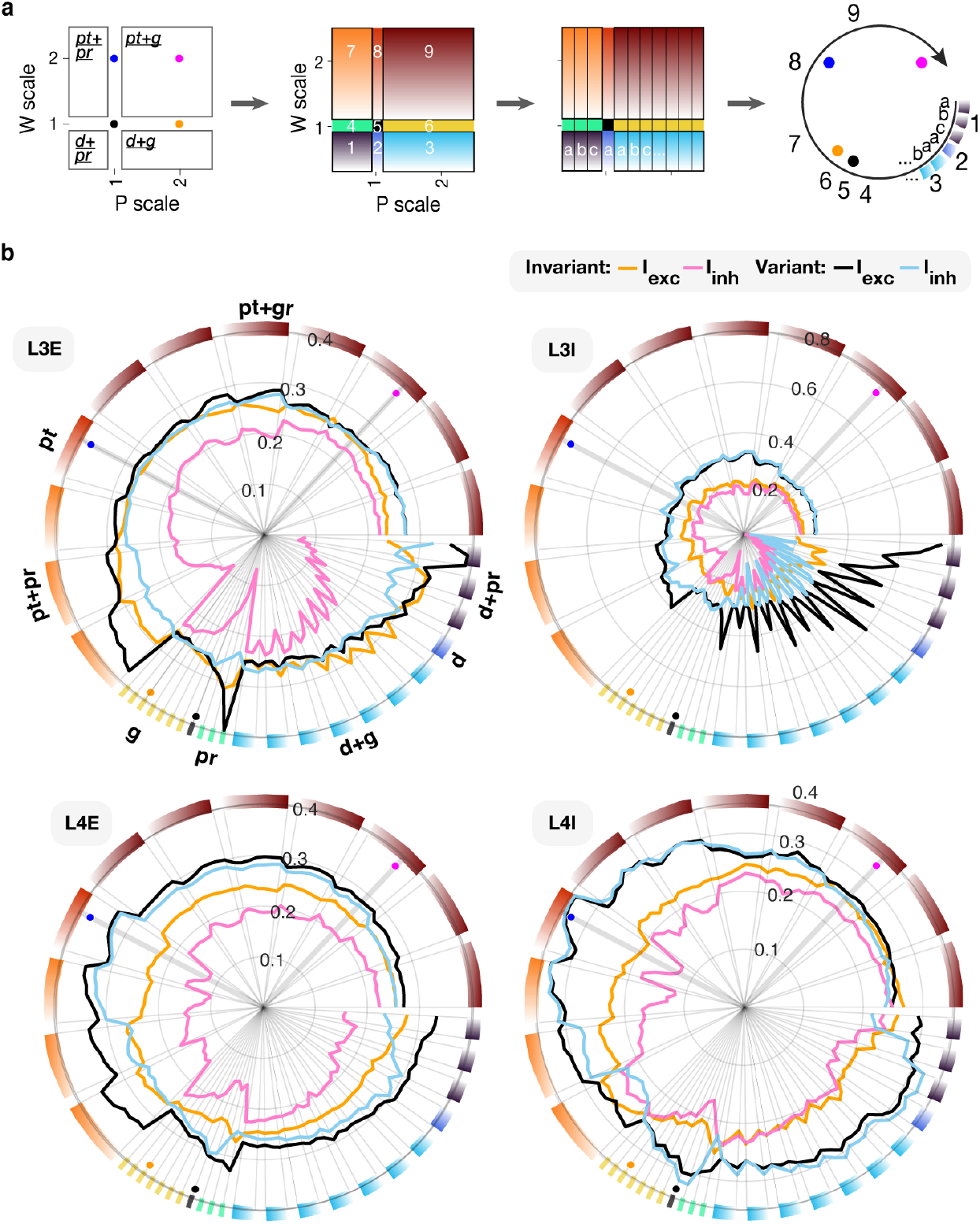
Connectivity-dependent modulation of excitatory and inhibitory presynaptic currents to input-invariant and variant neurons across four cortical populations. (a) The first cartoon shows a 2D matrix representing synaptic weight scaling (W, vertical axis) and connectivity probability scaling (P, horizontal axis), familiar from Figure 1c. The matrix is divided into four quadrants corresponding to combinations of connectivity scaling effects: potentiation (pt, W > 1), depression (d, W < 1), growth (g, P > 1), and pruning (pr, P < 1). Conditions between quadrants represent pure scaling effects, with only one type of change (pt, d, g, or pr). Dots indicate the four network conditions analysed in the main text: W1P1 (black), W1P2 (orange), W2P1 (blue), and W2P2 (pink). In the second cartoon, the same matrix is colour-coded into nine fields to distinguish quadrants and between-quadrant conditions. Colours are assigned arbitrarily but numbered sequentially for orientation, starting from the bottom-left quadrant and moving rightwards, then one row up, repeating left to right. These numbers provide a reference for the circular arrangement shown in the fourth cartoon. The third cartoon shows each colored field vertically subdivided into bars representing subgroups of conditions, labelled a, b, c… from left to right. Bars filled with a gradient indicate fixed P with increasing W, while solid-colour bars (fields 4-6) represent fixed P and W (see legend on top). In the fourth cartoon, the fields (numbers) and subgroups (letters) are arranged in a clockwise circle, starting with the quadrant of the lowest number at the 3 o’clock position, and continuing through letters and numbers clockwise. Colored dots mark the positions of the four analysed networks on the circle. (b) Each polar plot shows one cortical population (L2/3E, L2/3I, L4E, L4I; labelled at the top left of each circle). The circular axis represents different connectivity changes, where bars along the outer rim (circular axis) are coloured to represent within- or between-quadrant W-P conditions (see a for condition labels in quadrants): potentiation (*pt*, W>1), depression (*d*, W<1), growth (*g*, P>1), and pruning (*pr*, P<1), as well as combinations (e.g., *pt+pr, d+g*). Within each circle, the radial axis shows the fraction of total presynaptic current directed to each neuronal subpopulation. Currents are divided by type and neuron class: excitation to input-invariant neurons (orange), inhibition to input-invariant neurons (pink), excitation to variant neurons (black), and inhibition to variant neurons (light blue). (c) Dots along the outer rim indicate the four network conditions analysed in the main text: W1P1 (black), W1P2 (orange), W2P1 (blue), and W2P2 (pink).

